# MYRF is Essential for Epicardial Development

**DOI:** 10.1101/2025.10.17.682841

**Authors:** Elise V. Stanley, Chao Gao, Jason R. McCarthy, Maria I. Kontaridis, Tongbin Wu

**Affiliations:** Department of Biomedical Research and Translational Medicine, Masonic Medical Research Institute, 2150 Bleecker Street, Utica, NY, 13501, USA

**Author notes:** Correspondence: Address for Correspondence: Tongbin Wu, Ph.D., Department of Biomedical Research and Translational Medicine, Masonic Medical Research, Institute, 2150 Bleecker Street, Utica, NY, 13501, USA.

## Abstract

Myelin Regulatory Factor (MYRF) is a transcription factor previously known for myelination of neurons in the central nervous system. Accumulating evidence has recently implicated *MYRF* mutations in the pathogenesis of cardiac urogenital syndrome (CUGS), in which patients exhibit a range of cardiac abnormalities, including atrial septal defect (ASD), ventricular septal defect (VSD), and hypoplastic left heart syndrome (HLHS). However, the mechanisms by which MYRF deficiency leads to cardiac anomalies remain poorly defined. Moreover, in part due to the lack of cell-type specific loss-of-function models, it is unclear which cardiac cell types contribute to the defects observed in MYRF-related CUGS. To address these questions, we examined the expression pattern of MYRF in the heart and found that it is most highly expressed in epicardial cells (EPCs) among all cardiac cell types. Importantly, *Myrf* global knockout (KO) mice displayed impeded epicardial development, with markedly reduced epicardial coverage of the myocardial surface. Furthermore, specifically deleting MYRF in EPCs results in severely degenerated epicardium, dramatically reduced epicardial derived cells (EPDCs), and thin myocardial wall. Conversely, ablating MYRF in cardiomyocytes (CMs) did not lead to any overt cardiac phenotypes, indicating that MYRF is dispensable for the developing CMs. Taken together, our findings suggest that compromised function of MYRF in EPCs may contribute to the pathogenesis of MYRF-related CUGS. To our knowledge, this is the first study to link impaired epicardial development to this disease.

## MAIN TEXT

Myelin Regulatory Factor (MYRF) is a transcription factor previously known for myelination of neurons in the central nervous system.^1^ Recently, studies have linked *MYRF* mutations to cardiac urogenital syndrome (CUGS),^2^ in which patients exhibit a myriad of cardiac abnormalities, including atrial septal defect (ASD), ventricular septal defect (VSD), and hypoplastic left heart syndrome (HLHS). However, the mechanisms through which MYRF deficiency leads to these cardiac anomalies remain poorly defined. A earlier report implicates that MYRF may play a cell autonomous role in cardiomyocytes (CMs), as *Myrf* global knockout (KO) Medaka (Japanese rice fish) display ventricular wall hypoplasia.^3^ Yet, partly due to the lack of cell-type specific loss-of-function models, it remains unclear which cardiac cell types might contribute to the defects observed in MYRF-related CUGS. To address this gap in knowledge, we examined the expression pattern of MYRF in the mammalian heart by analyzing spatial transcriptomics data collected from E13.5 mouse hearts.^4^ We found that *Myrf* is enriched at peripheral portions of the heart (Figure [A]). To pinpoint the precise cell types in which MYRF is expressed, we performed MYRF immunofluorescence (IF) staining on E13.5 mouse hearts. The results show that MYRF is most highly expressed in the epicardial cells (EPCs) (Figure [B]), which is consistent with a scRNA-seq data in human fetal heart.^5^

**Figure 1.**
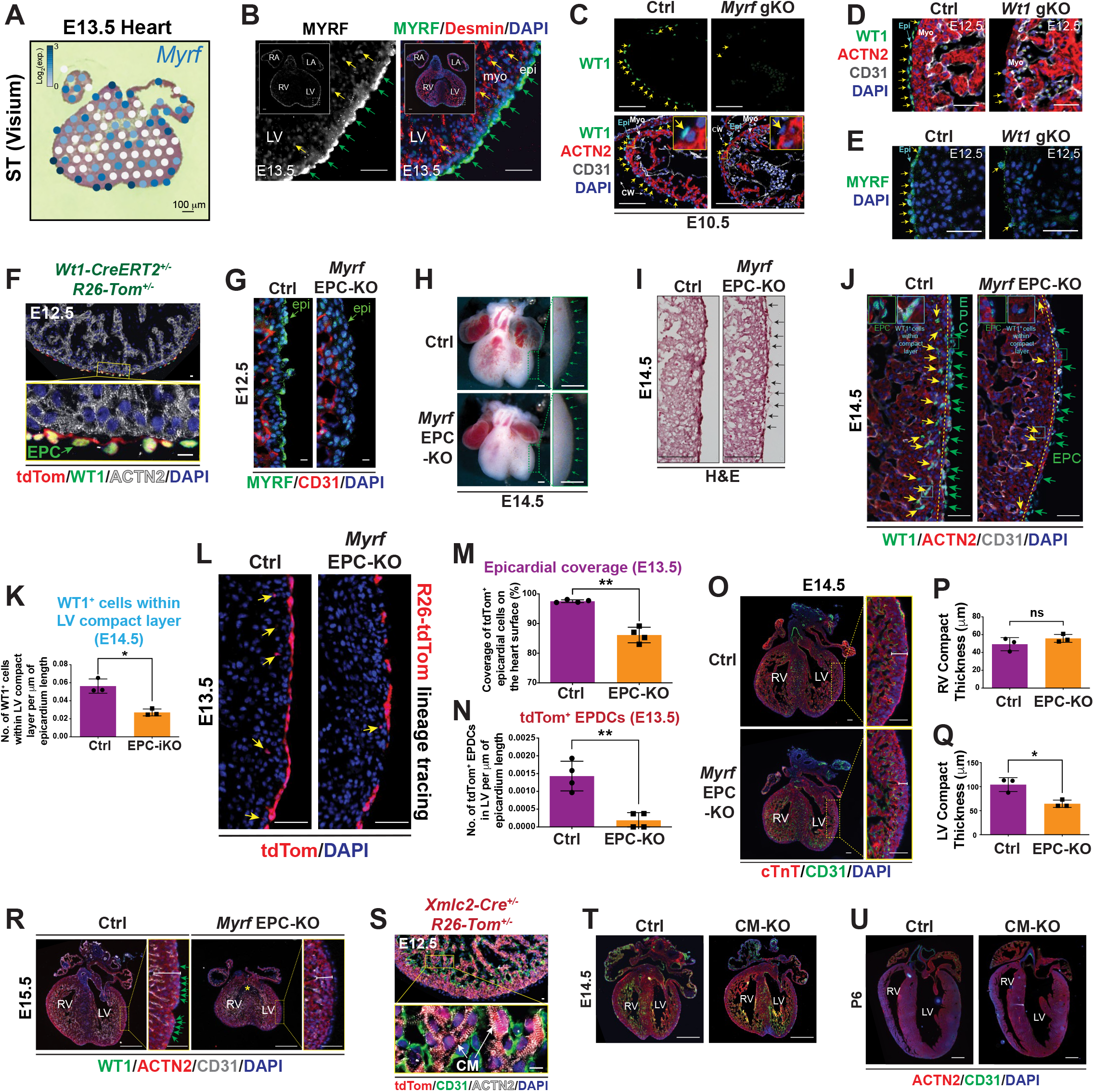
MYRF is essential for epicardial development. **A**, *Myrf* expression pattern visualized in Visium spatial transcriptomics (ST) data collected from embryonic day 13.5 (E13.5) mouse hearts. **B**, Representative immunofluorescence (IF) images of E13.5 mouse hearts stained with MYRF and desmin (DES) antibodies. epi: epicardium; myo: myocardium. LV: left ventricle. Green arrows indicate epicardial cells (EPCs); yellow arrows indicate cardiomyocytes (CMs). Scale bar: 50 μm. **C**, Representative IF images of E10.5 *Myrf* gKO and littermate control (ctrl) hearts. Yellow arrows indicate EPCs. WT1, EPC marker; ACTN2, α-actinin-2 (CM marker); CD31, endothelial cell (EC) marker. Scale bar, 100 μm. cw: chest wall tissue. Insets show WT1 expression in EPCs. **D-E**, Representative IF images of E12.5 *Wt1* gKO and littermate control hearts. Yellow arrows indicate EPCs. Scale bar, 50 μm. **F**, *Rosa26-tdTomato* lineage tracing of the *Wt1-CreERT2* line in E12.5 hearts (induced at E8.5). tdTom, tdTomato. Scale bar, 10 μm. **G**, MYRF expression in E12.5 *Myrf* EPC-KO and littermate control (*Myrf*^*flox/+*^*;Wt1-CreERT2*^*+/-*^) hearts. Scale bar, 10 μm. **H-I**, Morphology (H) and H&E staining (I) of E14.5 *Myrf* EPC-KO and control hearts. Arrows indicate partly detached epicardium. Scale bar, 100 μm. **J**, E14.5 *Myrf* EPC-KO and control hearts stained with antibodies as indicated. Green arrows indicate WT1^+^ EPCs. Yellow arrows indicate WT1^+^ cells within compact layer. Yellow dashed lines separate epicardium and myocardium. Scale bar: 50 μm. **K**, Quantification of WT1^+^ cells in LV compact layer of E14.5 *Myrf* EPC-KO and control hearts. n=3 per group. *p<0.05 (Welch’s t-test). **L-N**, R26-tdTom lineage tracing images (L) and quantification (M-N) of EPCs and EPDCs (yellow arrows) in E13.5 *Myrf* EPC-KO and control hearts. Scale bar: 50 μm. n=4 per group. **p<0.01 (Welch’s t-test). **O**, IF images of E14.5 *Myrf* EPC-KO and control hearts stained with antibodies as indicated. Scale bar: 100 μm. Rulers indicate compact myocardium thickness. **P-Q**, Quantification of RV and LV compact myocardium thickness of E14.5 *Myrf* EPC-KO and control hearts. n=3 per group. *p<0.05; ns, not significant (Welch’s t-test). **R**, IF images of E15.5 *Myrf* EPC-KO and control hearts stained with antibodies as indicated. Green arrows: WT1^+^ EPCs. Yellow asterisk indicates VSD in *Myrf* EPC-KO heart. Scale bar: 500 μm. **S**, *Rosa26-tdTomato* lineage tracing of the *Xmlc2-Cre* line in E12.5 hearts. Scale bar, 10 μm. **T-U**, E14.5 and postnatal day 6 (P6) *Myrf* CM-KO (*Myrf*^*flox/flox*^*;Xmlc2-Cre*^*+/-*^) and control hearts stained with antibodies as indicated. Scale bar: 500 μm.

To determine the systemic requirement of MYRF in heart development, we generated *Myrf* global KO (gKO) mice by crossing *Myrf* floxed mice^1^ with CMV-Cre. *Myrf* gKO mice died between E11.5 and E12.5 (data not shown). Strikingly, at E10.5, there were very few EPCs on the myocardial surface of *Myrf* gKO hearts (arrows, Figure [C]), and they did not express WT1, a transcription factor required for epicardial development^6^ (insets, Figure [C]). Interestingly, *Wt1* gKO mice (*Wt1-CreERT2*^*+/+*^) also had scarce EPCs, which expressed MYRF but not WT1 (Figures [D-E]), suggesting that the downregulation of WT1 may at least partially account for the epicardial phenotype in *Myrf* gKO mice. To determine whether the impaired epicardial development of *Myrf* gKO mice was due to the loss of cell autonomous function of MYRF in EPCs, we crossed *Myrf* floxed mice with inducible epicardial-specific *Wt1-CreERT2*^7^ mice to generate *Myrf* EPC-KO mice (*Myrf*^*flox/flox*^*;Wt1-CreERT2*^*+/-*^, induced at E8.5). The specificity of *Wt1-CreERT2* was verified by lineage tracing using *Rosa26-tdTomato* (R26-tdTom) indicator (Figure [F]). EPC-specific deletion of *Myrf* was confirmed by MYRF IF performed on E12.5 *Myrf* EPC-KO and control hearts (*Myrf*^*flox/+*^*;Wt1-CreERT2*^*+/-*^) (Figure [G]). Remarkably, at E14.5, the epicardial layer of EPC-KO heart was thinner than controls (green arrows, Figure [H]). In addition, histological analysis revealed that the epicardium partially detached from myocardium in EPC-KO hearts (Figure [I]), indicative of epicardial degeneration. To further confirm this, we performed IF with EPC marker WT1 using E14.5 EPC-KO and control hearts, and observed marked decrease of WT1^+^ EPCs and WT1^+^ cells within the LV compact myocardium of EPC-KO mice (Figures [J-K]). Notably, the expression level of WT1 in WT1^+^ cells appeared to be lower in EPC-KO (insets, Figure [J]), corroborating findings in the *Myrf* gKO mice.

EPCs give rise to epicardial-derived cells (EPDCs) through epithelial-to-mesenchymal transition (EMT), which then differentiate into several cardiac cell types.^8^ The reduction of WT1^+^ cells (may include EPDCs^9^) within the EPC-KO compact myocardium (Figures [J-K]) indicates that EPC EMT might be affected. To unequivocally demonstrate this, we performed R26-tdTom lineage tracing of EPCs (induced at E8.5 and analyzed at E13.5) and observed a pronounced reduction in epicardial coverage and tdTom^+^ EPDCs in EPC-KO hearts (Figures [L-N]). In line with the role of epicardium in supporting myocardial growth, the LV compact myocardium thickness was markedly reduced in EPC-KO while that of RV was not significantly affected (Figures [O-Q]). Eventually, most *Myrf* EPC-KO mice succumbed to death between E15.5 and E16.5, with the surviving mice retaining few WT1^+^ EPCs and having multiple morphological abnormalities, including thin compact layer and VSD, a phenotype also observed in MYRF-related CUGS^2^ (Figure [R]).

Although MYRF is expressed at a lower level in CMs than in EPCs (Figure [B]), it may still play a role in CMs. To test this, we generated *Myrf* CM-specific KO mice with *Xmlc2-Cre*.^4,10^ The specificity of *Xmlc2-Cre* was validated with R26-tdTom lineage tracing (Figure [S]). Surprisingly, these mice did not exhibit any apparent cardiac phenotypes (Figures [T-U]), indicating that MYRF does not play an important cell autonomous role in the developing CMs.

Collectively, our data suggest that MYRF is essential for epicardial development and ventricular wall morphogenesis. Importantly, compromised function of MYRF in EPCs may contribute to the pathogenesis of MYRF-related CUGS. To our knowledge, this is the first study to link impaired epicardial development to this disease.

## METHODS

### Mice

*Myrf* floxed mice were purchased from the Jackson Laboratory (stock no. 010607). All mice were of C57BL/6NCrl and C57BL/6J mixed background. Genotypes of mice were determined by PCR using *Myrf* flox primers (forward: 5’-GGGTACTAAAGAATGGCGAAGG- 3’, reverse: 5’- GTGGCAGGCACTAGAAGAGG -3’) and Cre primers (forward: 5’- GTTCGCAAGAACCTGATGGACA-3’, reverse: 5’-CTAGAGCCTGTTTTGCACGTTC-3’).

All animal procedures were performed in accordance with the National Institutes of Health Guide for the Care and Use of Laboratory Animals and approved by the Institutional Animal Care and Use Committees of the Masonic Medical Research Institute.

### Histology and immunofluorescence

Histology and immunofluorescence were performed as previously described.^4,10^ Briefly, mouse hearts were dissected and fixed with 4% paraformaldehyde (PFA) overnight at 4°C, incubated in 5%, 10%, 15%, 20% sucrose dissolved in PBS, embedded in OCT Tissue-Tek (Thermo Fisher Scientific), and then cut into 6 μm cryosections. For histology, sections were stained with hematoxylin and eosin (H&E) as previously described.^4^. For immunofluorescence, sections were blocked with PBST (1% BSA, 0.2% Tween-20 in PBS) for one hour at room temperature (R.T.), and incubated with primary antibody solution (antibodies diluted in PBST supplemented with 5% donkey serum) overnight in a humidified chamber at 4°C. Sections were then washed three times with PBST and then incubated with secondary antibody solution for 2 hours at R.T. Sections were counterstained with DAPI and mounted in Prolong Diamond Antifade mountant (Thermo Fisher Scientific). Images were captured using Keyence fluorescence microscope. Sources of antibodies used in immunofluorescence are: MYRF, BS-11191R (Bioss); Desmin, sc-7559 (Santa Cruz Biotechnology); WT1, 83535S (Cell Signaling); ACTN2, A7811 (Sigma-Aldrich); CD31, 550274 (BD Pharmingen); cTnT, MS295P1 (Fisher Scientific).

### Statistical Analysis and Data Availability

Data are presented as mean ± standard error of the mean (s.e.m.). Statistical analysis was performed using GraphPad Prism 10 software with Welch’s t test. *P*-values less than 0.05 were considered significant and reported as *p < 0.05, **p < 0.01, ***p < 0.001, ****p<0.0001. Detailed data, protocols, and materials can be obtained from the corresponding author upon reasonable request.

## ACKNOWLEDGEMENTS

We thank Chase Kessinger (Masonic Medical Research Institute) for technical assistance in immunofluorescence and histological studies.

